# Fixed-dose streptozotocin combined with high-fat diet induces a stable type 2 diabetes-like phenotype in C57BL/6J mice

**DOI:** 10.64898/2026.06.01.728717

**Authors:** Claes Fryklund, Christian Simonsson, Alexandra Hellberg, Josefin Malmberg, Karin G. Stenkula, Maria Swanberg

## Abstract

High-fat diet (HFD) combined with streptozotocin (STZ) is widely used to model type 2 diabetes (T2D) in rodents, but is often associated with high mortality, non-responders, and inconsistent outcomes. STZ is conventionally administered using body weight-adjusted dosing (mg/kg), despite evidence that heavier animals, including HFD-fed mice, exhibit more severe glycaemic responses.

Here, we performed metabolic phenotyping in chow- and HFD-fed C57BL/6J mice treated with low or high fixed doses (mg instead of mg/kg) of anomer-equilibrated STZ. HFD combined with low-dose STZ induced a stable T2D-like phenotype characterized by sustained obesity, moderate hyperglycaemia, insulin resistance, and partial β-cell loss, with low inter-individual variability. In contrast, high-dose STZ induced a T1D-like phenotype with extensive β-cell loss. A semi-mechanistic mathematical model was developed and validated against independent experimental data, reproducing the observed dynamics of fasting glucose in response to fixed-dose STZ. The model further predicted that weight-adjusted (mg/kg) dosing could introduce variability in glycaemic responses, particularly in HFD-fed mice.

Together, these results demonstrate that fixed-dose, anomer-equilibrated STZ induces a stable T2D-like phenotype, providing an alternative to conventional weight-adjusted dosing in HFD-fed mice.

## 1. Introduction

Type 2 diabetes (T2D) is a major global health problem, with recent estimates indicating that approximately 500-800 million people are affected (1, 2). The disease substantially increases the risk of cardiovascular complications, including stroke and coronary heart disease, and was ranked as the eighth leading cause of death globally in 2019 (3, 4). T2D is a complex, polygenic disease arising from the interplay of genetic, epigenetic, and environmental factors, with lifestyle factors such as physical activity and energy intake playing central roles (5). Its pathogenesis is heterogeneous and involves impairments in pathways related to β-cell function, insulin action, incretin signalling, and adipose tissue distribution (6). This heterogeneity has motivated the use of experimental mouse models that capture distinct aspects and stages of T2D pathology, from early insulin resistance and hyperinsulinemia (prediabetes) to late-stage β-cell dysfunction and hyperglycaemia. Among the most commonly used models are diet-induced obesity via high-fat diet (HFD), genetically modified strains (e.g. *ob/ob*, *db/db*), and toxin-induced β-cell depletion using streptozotocin (STZ) (7).

C57BL/6J mice fed HFD exhibit insulin resistance, obesity, and mild hyperglycaemia, representing a prediabetic state of T2D (8). These mice resist progression to overt T2D by maintaining near-normal glucose through compensatory β-cell expansion and enhanced insulin secretion (9, 10). To induce overt T2D in C57BL/6J mice, the β-cell toxin STZ is often administered in combination with HFD feeding (11). STZ is taken up by pancreatic β-cells via GLUT2 transporters, where it causes DNA damage and β-cell death, leading to sustained hyperglycaemia (12).

The combination of repeated low-dose STZ injections after a few to several weeks of HFD feeding is a common approach to model T2D. However, there is no fully standardized protocol, and large variations in diet duration, STZ dosage, and injection frequency likely contribute to inconsistent outcomes, including high mortality and non-responsive animals (13). Moreover, variability in response to STZ may arise from different protocols for STZ preparation, leading to differences in α/β-anomeric composition. As soon as STZ is dissolved, the more toxic α-anomer gradually converts to the less toxic β-anomer, a process that requires at least two hours to reach an α/β-anomeric equilibrium (14). Immediate injection of STZ after dissolution has long been recommended practice due to presumed instability, but studies have demonstrated that STZ remains stable and biologically active for several days when stored cold and dark in acidic buffer (14, 15). Recent studies recommend α/β-anomer equilibration (≥ 2h) for improved reproducibility and reduced mortality (14, 16).

One largely unexplored area in STZ-based models concerns the dosing strategy, as STZ has historically been administered using body weight-adjusted (mg/kg) dosing rather than fixed dosing (mg). Previous studies in rodents have shown that body weight-adjusted dosing results in more severe hyperglycaemia in heavier animals (11, 17). HFD-fed mice present a particular challenge due to their highly variable weight gain, typically ranging ∼10-50 % above chow controls. STZ is hydrophilic and mainly distributes in lean body mass, not adipose tissue (18, 19). Since HFD-induced weight gain comes from fat accumulation, whereas β-cell toxicity from STZ most likely depends on lean mass, body weight-adjusted dosing risks overdosing obese mice. This can potentially cause increased mortality in heavier animals, and, in contrast, non-responding lean mice. We therefore hypothesized that fixed (non-weight-adjusted) STZ dosing would lead to more uniform β-cell toxicity across HFD-fed mice regardless of individual weight gain, improving reproducibility while minimizing non-responders and mortality. To test this hypothesis, we examined whether HFD combined with fixed, anomer-equilibrated STZ would induce a stable T2D-like phenotype, using longitudinal measures of glucose homeostasis, insulin resistance, β-cell function and pancreatic histology. The results were integrated in a systems biology mathematical model that predicted diet, STZ, and time-dependent responses, and was validated in independent *in vivo* data.

## 2. Material and Methods

### 2.1 Animals and study outline

C57BL/6J mice were purchased from Taconic and maintained by in-house breeding. Male offspring mice were housed in groups of 4–5 mice per cage, with ad libitum access to food and water. At 8 weeks of age (study week 0), mice were assigned to either an HFD (60% kcal from fat, #D12492, Research Diets) (n = 19) or chow control diet (14% kcal from fat, #SAFE A30, SAFE Diets, France) (n = 20). At week 7, mice received STZ (see details below) or vehicle for 4–5 consecutive days (see Fig. 1 for study outline). The experimental groups were: chow vehicle (n = 10), HFD vehicle (n = 7), chow-STZ^low^ (n = 5), HFD-STZ^low^ (n = 7), chow-STZ^high^ (n = 5), and HFD-STZ^high^ (n = 5). The study endpoint was week 25, and body weight was measured biweekly throughout the study. One mouse in each of the HFD-STZ^high^ and chow-STZ^high^ groups was euthanized due to blood glucose ≥30 mM (humane endpoint). One mouse in the chow-STZ^low^ group died during handling.

**Figure 1.**
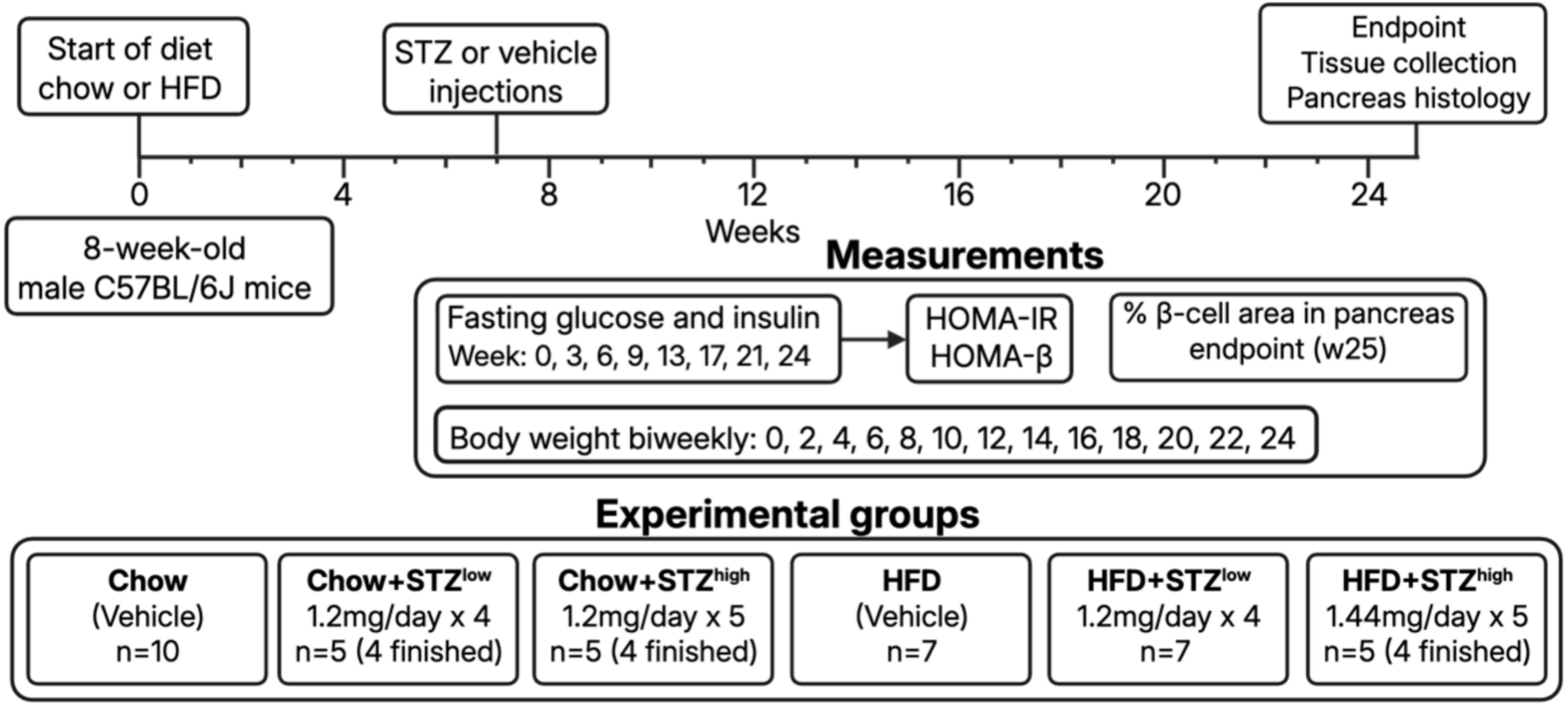
Study outline and experimental timeline. Eight-week-old male C57BL/6J mice were assigned to either chow or high-fat diet (HFD) starting at week 0. At week 7, mice received daily intraperitoneal injections of streptozotocin (STZ) or vehicle for 4–5 consecutive days to induce β-cell dysfunction. Metabolic parameters, including fasting blood glucose and insulin, were measured at weeks 0, 3, 6, 9, 13, 17, 21, 24 and used to calculate HOMA-IR and HOMA-β. Body weight was recorded biweekly throughout the study. At the endpoint (week 25), tissues were collected and pancreatic β-cell area was assessed histologically. Experimental groups included: chow (n = 10), chow+STZ^low^ (1.2 mg/day × 4; n = 5, n = 4 completed), chow+STZ^high^ (1.2 mg/day × 5; n = 5, n = 4 completed), HFD (n = 7), HFD+STZ^low^ (1.2 mg/day × 4; n = 7), and HFD+STZ^high^ (1.44 mg/day × 5; n = 5, n = 4 completed).

### 2.2 Streptozotocin treatment

STZ (≥75% α-anomer basis, S0130-100, Sigma) was dissolved in sterile filtered (Filtropur S, 0.2 µm, Sarstedt) sodium citrate buffer (0.05 M, pH 4.5) ≥2 h prior to the first injection to ensure equilibrium (44:56) between α- and β-anomers (14). The vial with STZ was kept dark at 4 °C and used for subsequent days (4-5 days). At week 7, STZ was administered by intraperitoneal injection, vehicle mice received sodium citrate buffer. Non-fasted mice (20) received low dose STZ during four consecutive days (STZ^low^: 1.2mg/day × 4 days; total 4.8mg), or high dose STZ during five consecutive days (chow-STZ^high^: 1.2mg/day × 5 days; total 6mg, HFD-STZ^high^: 1.44mg/day × 5 days; total 7.2mg). Mice receiving STZ were supplemented with 10% sucrose in their drinking water for 7 days post injection to prevent hypoglycaemia.

### 2.3 Tissue collection and assessment of pancreatic β-cell area

At experimental endpoint, mice were euthanized by intraperitoneal injection of pentobarbital (100-200 mg/kg), followed by transcardial perfusion with saline, followed by 4% paraformaldehyde (PFA). Epididymal and inguinal fat depots were dissected and weighed. Pancreas was dissected, post-fixed in 4% PFA overnight at 4°C, dehydrated in 70% ethanol, and paraffin-embedded. Sections were cut using a Leica RM2265 microtome. Blocks were trimmed until the complete pancreas outline was visible (0 µm), and 4-6 sections of 4 µm thickness were collected at +0, +400, and +800 µm intervals. Sections were transferred to slides (Superfrost Plus) and then deparaffinized in xylene, rehydrated through graded ethanol, and subjected to antigen retrieval (10 mM sodium citrate, pH 6.0, 40 min, 95°C) followed by quenching of endogenous peroxidase activity (3% H₂O₂, 10 min). Samples were blocked in 5% goat serum in 0.25 % PBST and incubated overnight at 4°C with a rabbit anti-insulin primary antibody (1:1000, Cell Signaling #3014T) in the same blocking buffer. Secondary staining was performed with biotinylated goat anti-rabbit (1:200, 1 h, RT, Vector laboratories #BA-1000), followed by ABC-HRP (Avidin/Biotin complex, Vector laboratories, #PK-6100, 30min, RT) and DAB chromogen (3,3’-diaminobenzidine, Vector laboratories #SK-4100, 5min, RT) according to instructions by manufacturer. Slides were counterstained with hematoxylin, dehydrated, and coverslipped using Pertex (Histolab, Sweden). Whole-slide images at 20x magnification from pancreas sections collected at +400 µm and +800 µm, were acquired using an Olympus BX61 with VS120-L100 automated stage (Olympus, Japan). Images were analysed blindly, using QuPath (21). The percentage of β-cell area was quantified by measuring insulin immunopositive area over total pancreatic area via haematoxylin and DAB channel thresholding, unspecific background staining outside the islets was manually excluded.

### 2.4 Blood sampling and calculation of HOMA-IR and HOMA-β

Fasting blood glucose and serum insulin levels were measured in mice following a 4-hour fast (7–11 a.m.). Blood was collected from the lateral saphenous vein, and glucose levels were determined using a handheld glucometer (Contour XT, Bayer). Approximately 20–30 µL of blood was collected into serum/clotting activator tubes (Microvette® CB 300, Sarstedt) and allowed to coagulate (15 min, RT). Samples were then centrifuged (1500 × g, 10 min, 4 °C) and the resulting serum was collected and stored at −80 °C. Serum insulin concentrations were quantified using an ELISA kit (Crystal Chem, #90080) according to the manufacturer’s instructions. Indices of insulin resistance (IR) and β-cell function (β) were calculated using the homeostatic model assessment (HOMA) method as previously described (22). HOMA-IR was calculated as insulin (µU/mL) × glucose (mM) / 22.5, and HOMA-β as (20 × insulin (µU/mL)) / [glucose (mM) − 3.5].

### 2.5 Mathematical modelling

The mathematical model was constructed using ordinary differential equations with all formal analysis performed in MATLAB 2023b (The MathWorks Inc., Natick, Massachusetts) using the systems biology toolbox (23). The full list of assumptions and model equations can be found in the supplementary material (Supplementary S3). The model parameter estimation was performed using the MEIGO toolbox in MATLAB (24).

### 2.6 Data and statistical analysis

GraphPad Prism v.10.6.1 (GraphPad Software) was used for statistical analyses. Pre-STZ baseline variables at week 6 were compared between chow and HFD groups using unpaired two-tailed Student’s t-test. A two-way ANOVA was performed on data post-STZ (average across weeks 9-24), with factors diet (chow, HFD) and STZ dose (vehicle, low, high), followed by Sidak’s post hoc multiple comparisons test. Assumptions of normal distribution and homoscedasticity of residuals were evaluated using the Shapiro–Wilk test, Q–Q plots, and residuals-versus-predicted plots. Variables that did not meet these assumptions were log2-transformed and re-analysed. Fasting insulin, HOMA-IR, HOMA-β, and percentage of β-cells required log2 transformation and were therefore analysed using two-way ANOVA on log2-transformed data.

## 3. Results

### 3.1 HFD combined with a fixed STZ dose mimics T2D pathology with sustained obesity and moderate hyperglycaemia

Body weight was monitored since HFD induces weight gain and STZ is known to cause weight loss. Before STZ treatment (week 6), the HFD groups had, as expected, significantly higher body weights compared to chow (p < 0.001, unpaired t-test) (Fig. 2A,B). At study endpoint (week 25) there was a significant main effect of STZ and diet, and a significant interaction of diet and STZ on body weight (p < 0.001). Post hoc analysis revealed that HFD-STZ^low^ was the only STZ-treated group that continued to gain weight and remained obese, ∼30% heavier compared to control at study endpoint (Fig. 2A,B). All other STZ-treated groups, including HFD-STZ^high^, stabilized at body weights slightly below those of chow-fed controls. Likewise, adiposity, measured as epididymal and inguinal depot weight, was significantly increased only in the HFD and HFD-STZ^low^ groups (Supplementary Fig. S1A,B).

**Figure 2.**
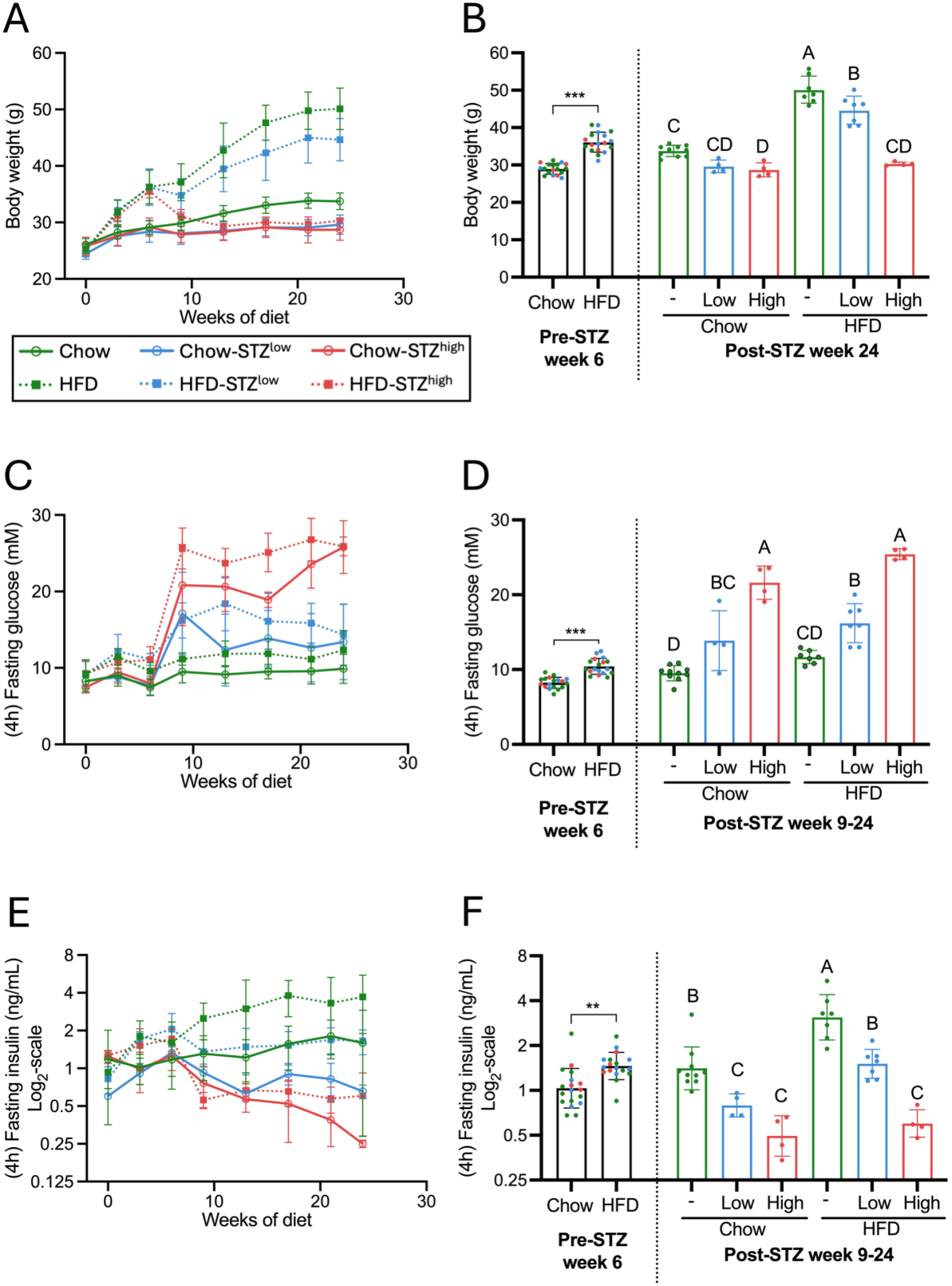
Fixed-dose anomer-equilibrated STZ combined with HFD induces stable hyperglycaemia while preserving weight gain in HFD-STZ^low^ mice. (A, C, E) Body weight, fasting blood glucose and fasting insulin were measured in chow- and HFD-fed mice receiving vehicle, STZ^low^, or STZ^high^ every 3–4 weeks over 24 weeks (body weight was measured biweekly). (B) Body weight at week 6 (pre-STZ) and week 24 (post-STZ), with the dashed line separating pre- and post-STZ data. (D) Fasting blood glucose at week 6 (pre-STZ) and the mean value from weeks 9–24 (post-STZ), with the dashed line separating pre- and post-STZ data. (F) Fasting insulin at week 6 (pre-STZ) and the mean value from weeks 9–24 (post-STZ), with the dashed line separating pre- and post-STZ data. Pre-STZ differences between chow and HFD were analysed by unpaired Student’s t-test, *p < 0.05, **p < 0.01, ***p < 0.001. Post-STZ differences were analysed by two-way ANOVA with diet and STZ treatment as factors, followed by Sidak’s multiple-comparisons test. Different letters indicate significant differences between groups based on Sidak’s post hoc multiple-comparisons test (p<0.05), groups sharing a letter are not significantly different. Data are shown as mean ± SD. Data from insulin (E,F) were log2-transformed for statistical analysis and are shown on a log2-transformed axis.

Fasting blood glucose levels were measured every 3-4 weeks. At week 6 (pre-STZ), HFD-fed mice exhibited a significant, mild elevation of glucose levels compared to chow-fed controls (p < 0.001, unpaired t-test) and this diet effect remained stable throughout the study (Fig. 2C,D). Post-STZ (weeks 9–24), two-way ANOVA revealed significant effects of diet and STZ dose on fasting blood glucose (p < 0.001), but no interaction (Fig. 2D). Sidak’s post hoc confirmed hyperglycaemia in all STZ-treated groups relative to chow controls, with STZ^high^ causing greater elevations than STZ^low^ (Fig. 2D). To evaluate whether a fixed-dose STZ strategy influenced blood glucose differently from conventional weight-adjusted dosing, the relationship between pre-STZ body weight and post-STZ fasting blood glucose was examined in chow-STZ^low^ and HFD-STZ^low^. Despite substantial variation in pre-STZ body weight (26.5–40.7 g), simple linear regression showed no association between body weight and post-STZ fasting blood glucose with fixed-dose STZ (slope= 0.23, 95% CI [-0.14 to 0.97], R^2^=0.16, p=0.21) (Supplementary Fig. S1C). The positive, though non-significant, slope highlights a potential concern that weight-adjusted dosing could result in overdosing heavier mice, since these mice would receive a larger absolute STZ dose under a weight-adjusted scheme.

### 3.2 HFD induces hyperinsulinemia while low-dose STZ normalizes insulin

In C57BL/6J mice, HFD induces proliferation of β-cells and hyperinsulinemia to compensate for insulin resistance, while STZ treatment causes β-cell degeneration and hypoinsulinaemia (9). We therefore monitored fasting insulin throughout the study to assess the effects of diet and STZ dose on insulin levels. HFD caused a significant increase in fasting insulin levels compared to chow-fed mice at week 6 (pre-STZ) (p < 0.01, unpaired t-test) (Fig. 2E,F). Post-STZ (w9-24), there was a significant main effect of diet and STZ (p < 0.001), but no interaction (Fig. 2F). Post hoc analysis showed that the HFD group had significantly higher insulin levels than all other groups and that STZ reduced insulin levels in a dose-dependent manner (Fig. 2F). Insulin levels in HFD-STZ^low^ mice remained comparable to chow controls and were significantly lower than in the HFD group (Fig. 2F). This partial reduction in insulin levels in HFD-STZ^low^ mice was sufficient to induce hyperglycaemia without causing weight loss, indicating an optimal balance between insulin levels and obesity for a sustained T2D-like phenotype.

### 3.3 HFD combined with low-dose STZ induces T2D hallmarks with both increased insulin resistance (HOMA-IR) and β-cell dysfunction (HOMA-β)

HOMA-IR and HOMA-β, calculated from fasting glucose and insulin correlate well with clamp-based measures, and were used as surrogate indices of insulin resistance and β-cell function, respectively (25). Pre-STZ (w6), HFD-fed mice had significantly increased insulin resistance (higher HOMA-IR) compared to chow-fed mice (p < 0.001, unpaired t-test) but no significant difference in β-cell function (HOMA-β) (Fig. 3A-D). Post-STZ treatment (w9-24), HOMA-IR showed significant main effects of diet and STZ dose (p < 0.001) as well as significant diet × STZ interaction (p < 0.05) (Fig. 3B). Post hoc analysis revealed that HFD and HFD-STZ^low^ were the only groups with significantly increased HOMA-IR compared to chow controls (Fig. 3B). For HOMA-β there was a significant main effect of STZ dose (p<0.001), but no effect of diet or interaction (Fig. 3D). Post hoc analysis showed that all STZ-treated groups exhibited marked reductions in HOMA-β compared to vehicle-treated groups, with a significantly greater reduction in STZ^high^ groups (Fig. 3D). Assessment of the relationship between HOMA-IR and HOMA-β across individual mice revealed clear clustering of metabolic phenotypes according to diet and STZ treatment (Supplementary Fig. S2A). Relative to chow controls, HFD mice clustered at high HOMA-IR together with elevated HOMA-β values. HFD-STZ^low^ mice formed a distinct cluster characterized by moderately increased HOMA-IR and reduced HOMA-β. In contrast, STZ^high^ mice clustered at normal HOMA-IR and low HOMA-β values, irrespective of diet (Supplementary Fig. S2A).

**Figure 3.**
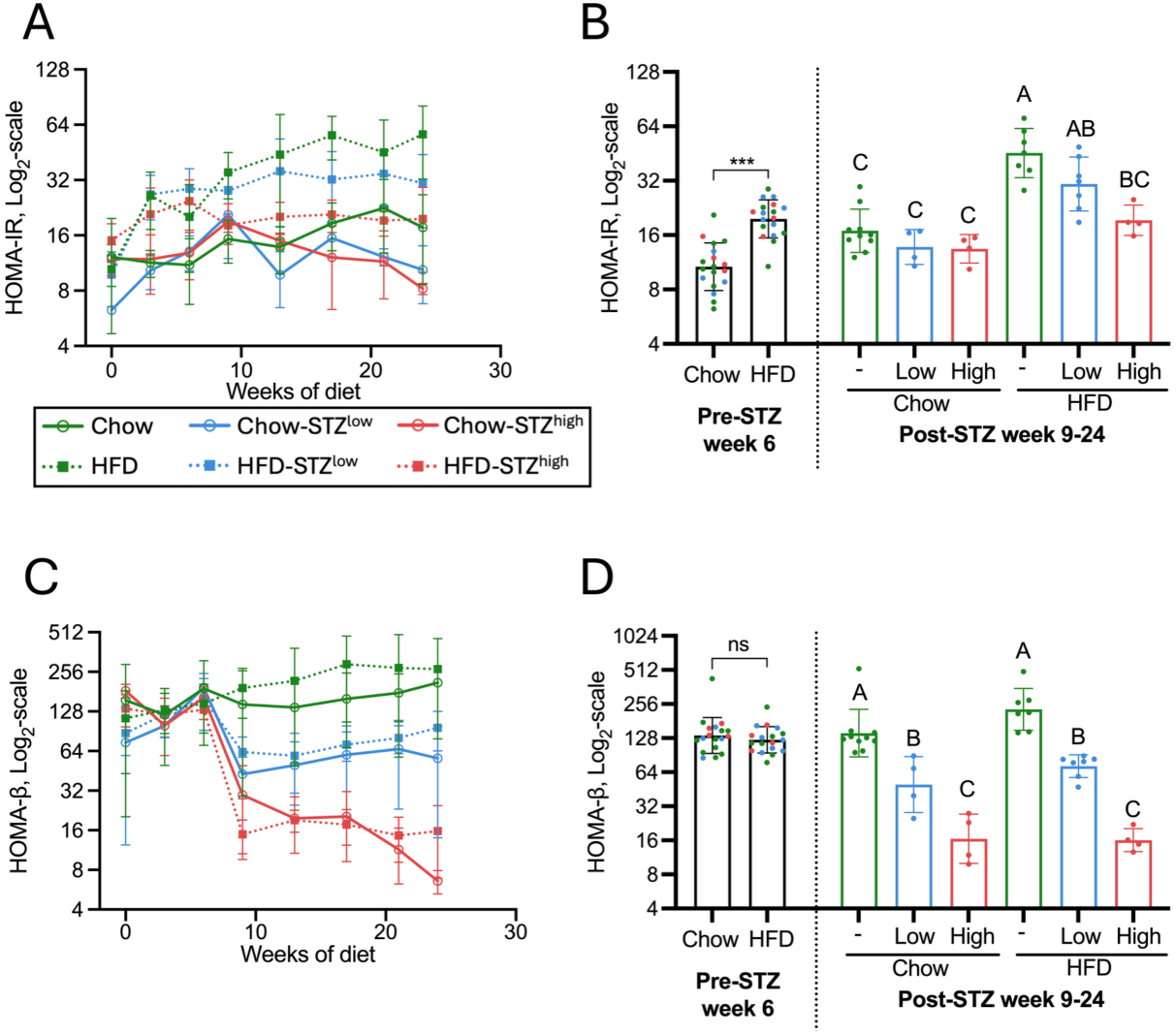
Indices of β-cell function (HOMA-β) and insulin resistance (HOMA-IR) reveal a robust T2D phenotype in HFD-STZ^low^ mice. (A, C) HOMA-IR and HOMA-β were measured in chow- and HFD-fed mice receiving vehicle, STZ^low^, or STZ^high^ every 3–4 weeks over 24 weeks. (B, D) HOMA-IR and HOMA-β at week 6 (pre-STZ) and the mean values from weeks 9–24 (post-STZ), with the dashed line separating pre- and post-STZ data Pre-STZ differences between chow and HFD were compared by unpaired Student’s t-test, *p < 0.05, **p < 0.01, ***p < 0.001. Post-STZ differences were analysed by two-way ANOVA with diet and STZ treatment as factors, followed by Sidak’s multiple-comparisons test. Different letters indicate significant differences between groups based on Sidak’s post hoc multiple-comparisons test (p<0.05), groups sharing a letter are not significantly different. Data were log2-transformed for statistical analysis and are shown on a log2-transformed axis as mean ± SD.

### 3.4 HOMA-β is a valid surrogate for β-cell loss after STZ treatment

At endpoint, β-cell loss was assessed by immunohistochemistry and quantification of insulin-positive staining in pancreas. There was a significant main effect of STZ dose (p < 0.001), but no significant effect of diet or interaction, and post hoc analysis revealed a STZ dose-dependent reduction in insulin-positive staining (Fig. 4A). The HFD-STZ^low^ group displayed significantly higher β-cell area than the HFD-STZ^high^ and chow-STZ^high^ groups, although still ∼50% lower than in normal, vehicle-treated groups. Simple linear regression on log₂-transformed data showed that HOMA-β strongly predicted the percentage of insulin-positive β-cell area (slope = 0.69, 95% CI [0.59–0.79], R^2^=0.85 p < 0.001) indicating that HOMA-β is a good predictor of STZ-induced β-cell loss (Fig. 4B).

**Figure 4.**
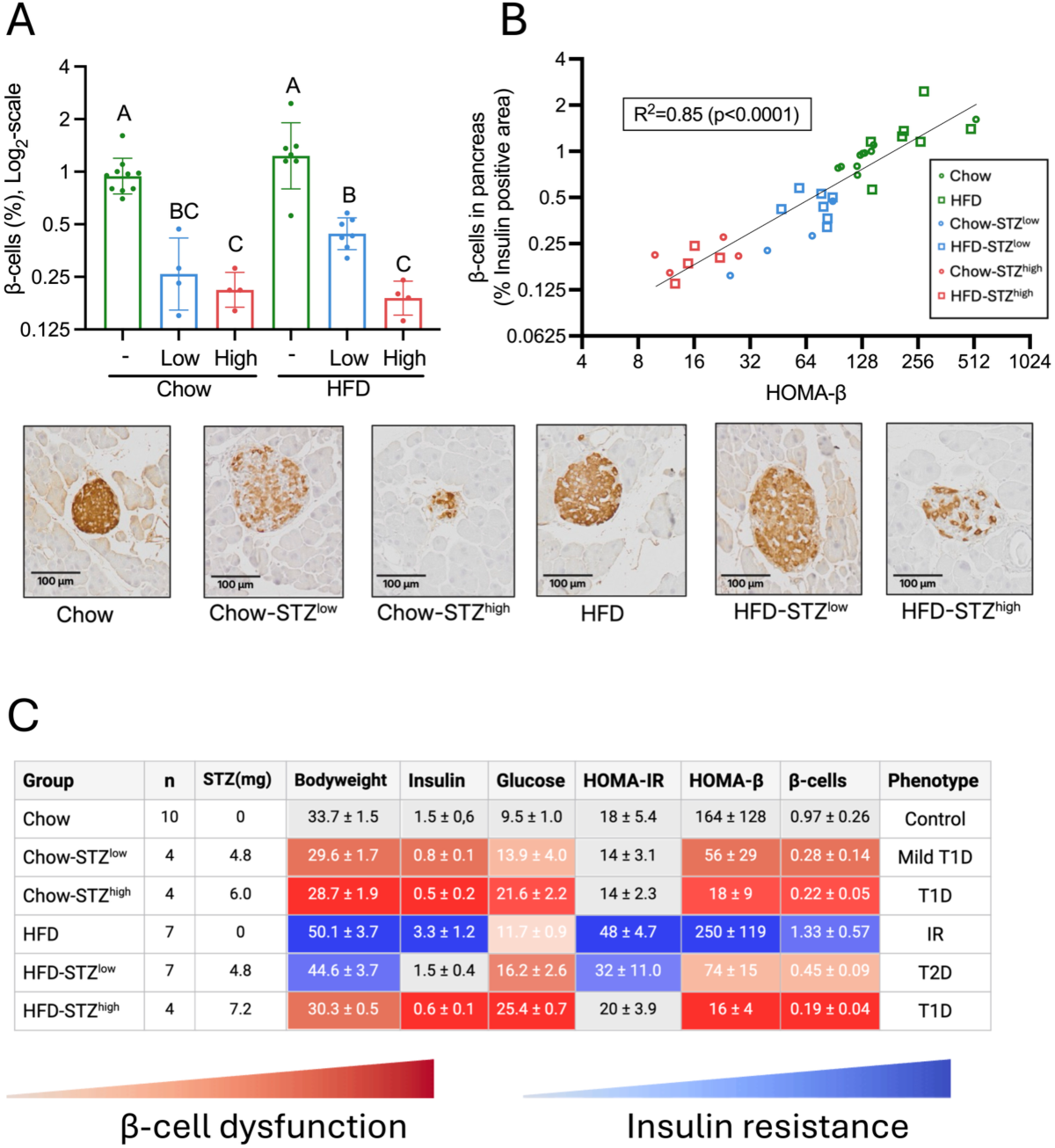
HOMA-β predicts insulin-positive β-cell area after STZ treatment. (A) Insulin-positive β-cell area, expressed as % of total pancreas area, was quantified at endpoint in chow-and HFD-fed mice receiving vehicle, STZ^low^, or STZ^high^. Data analysed by two-way ANOVA with diet and STZ treatment as factors, followed by Sidak’s multiple-comparisons test. Different letters indicate significant differences between groups based on Sidak’s post hoc multiple-comparisons test (p<0.05), groups sharing a letter are not significantly different. Data were log2-transformed for statistical analysis and are shown on a log2-transformed axis as mean ± SD. (B) Linear regression analysis of log2-transformed HOMA-β versus insulin-positive β-cell area showed a strong positive correlation (R2=0.85, slope=0.69, 95% CI 0.59–0.79, p<0.0001). Representative insulin-stained pancreatic sections are shown below. (C) Summary and overview of metabolic phenotypes across groups based on body weight, fasting insulin and glucose, HOMA-IR, HOMA-β, and β-cell quantification (insulin-positive area/total pancreas area). Values represent group mean ± SD calculated from the average value of each mouse across weeks 9–24 post-STZ treatment, except for body weight and β-cell measurements, which were assessed at endpoint, body weight at week 24 and β-cell area at terminal collection (week 25). Red and blue colour gradients indicate increasing β-cell dysfunction and insulin resistance, respectively, while grey indicates metabolically healthy reference values (chow). Note that fasting glucose reflects both insulin resistance and β-cell dysfunction but is represented here using red gradient (β-cell dysfunction).

### 3.5 Combination of multiple metabolic parameters across intervention groups identifies distinct models for insulin resistance, T1D, and T2D

The six experimental groups allowed the combined investigation of diet- and STZ-induced effects on metabolic phenotypes (Fig. 4C). The chow vehicle group represented metabolically healthy controls and served as a reference for metabolic changes in the other groups. The HFD vehicle group displayed a phenotype characterized by insulin resistance, consistent with a prediabetic state maintained by compensatory β-cell function. The chow-STZ^low^ group exhibited mild β-cell dysfunction, consistent with early/mild T1D. Both STZ^high^ groups (chow and HFD) displayed metabolic profiles associated with severe β-cell dysfunction and hyperglycaemia, consistent with overt T1D. The HFD-STZ^low^ group was the only group exhibiting both β-cell dysfunction and insulin resistance, consistent with a T2D-like phenotype. Thus, by different combinations of diet and STZ dosing, we found distinct models for insulin resistance (prediabetes), T1D (mild/overt), and T2D in C57BL/6J mice. Additionally, the HFD-STZ^low^ group demonstrated low inter-individual variability across key T2D parameters (averaged across weeks 9-24): blood glucose (13-20 mM), HOMA-IR (18–49), HOMA-β (47–88), β-cell area (0.44–0.58%), and body weight (40.4–50.1 g) (min-max range). This finding indicates that fixed-dose STZ strategy produced a stable T2D-like phenotype with low inter-individual variability.

### 3.6 Mathematical modelling of β-cell loss after STZ treatment

We created a semi-mechanistic mathematical model to further elucidate the relationship between the STZ dosage and glucose response. The model consisted of ordinary differential equations to describe the β-cell, glucose and insulin dynamics after STZ treatment (Fig. 5A). The full model description can be found in the supplementary (Supplementary S3). The model was trained to simultaneously describe collected data at week 0, 6, 9 and 24 for β-cell, glucose and insulin; control, STZ^low^ and STZ^high^, both for chow (Fig. 5B) and HFD (Fig. 5C). This resulted in a model behaviour representing the mean dynamics in all data. Overall, the model fits qualitatively well to all data. The best model agreement can be seen as the line, and the model uncertainty (best agreement + χ^2^(0.95,1)) is the shaded area. Next the predictive capabilities of the model were validated using independent-validation fasting plasma glucose data from Gilbert et al. (11) from mice undergoing STZ treatment with body weight-adjusted dose. The model was able to qualitatively well predict the levels of plasma glucose for both the chow and HFD (Fig. 5D).

**Figure 5.**
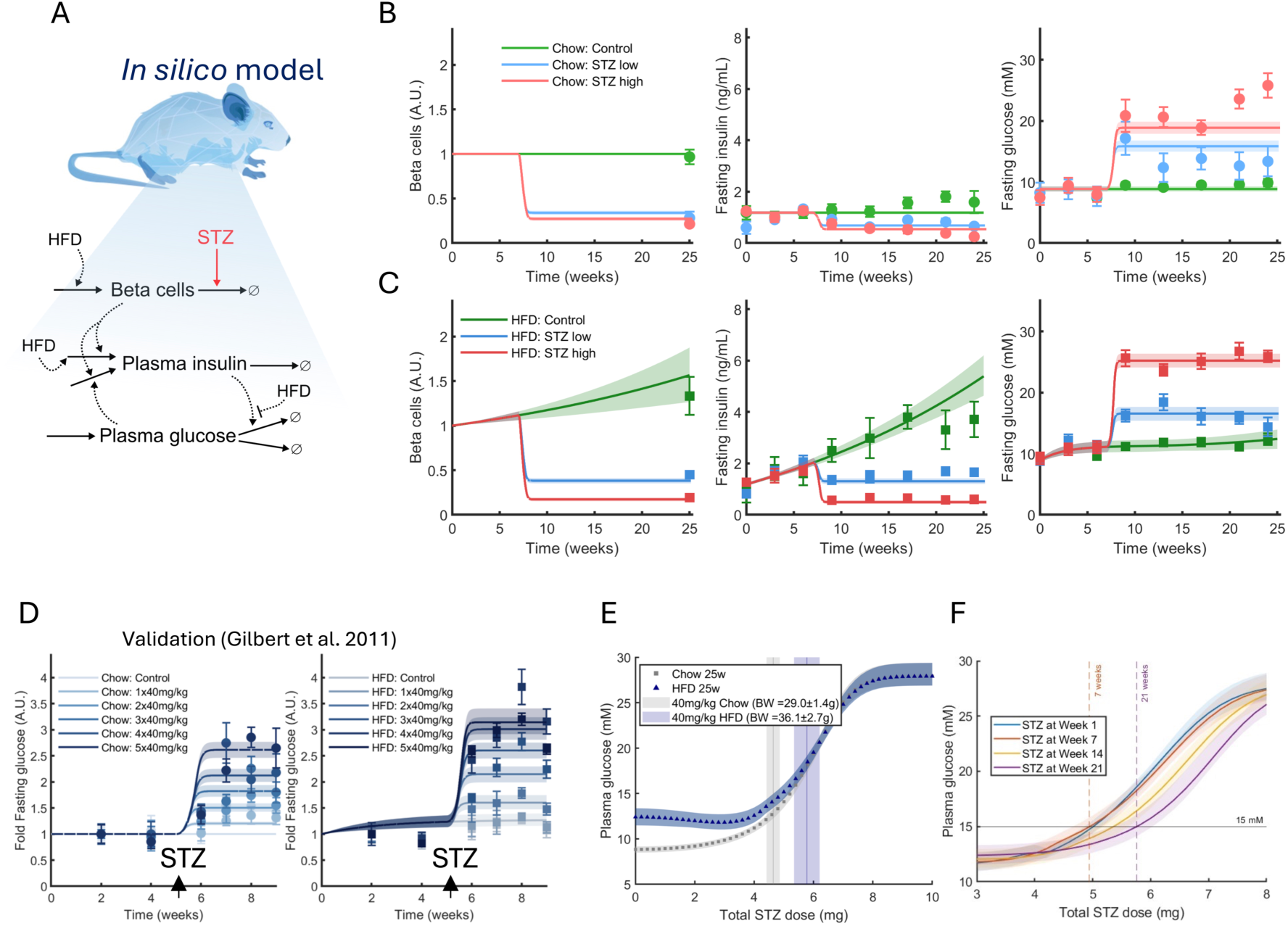
Mathematical Modelling of STZ intervention in mice. (A) Schematic representation of the mathematical model structure. (B,C) Simultaneous agreement between model simulations and all estimation data. Here, the model simulations (lines) for vehicle (green), STZ^low^ (blue), or STZ^high^ (red) for chow (B) and HFD (C) are compared to the experimental data (error bars), for β-cells, fasting insulin and fasting plasma glucose. (D) Model validation using independent experimental data from Gilbert et al. Model predictions of fasting plasma glucose for STZ dosing schemes not included during model fitting were compared with independent experimental data. Solid lines represent model predictions, while points and error bars show the corresponding experimental data. (E) Model predictions of fasting plasma glucose at 25 weeks when administering different levels of total STZ doses for chow and HFD. The dots are the endpoint of each 25 week in silico trial, chow is grey and HFD is blue, with the surrounding area indicating model uncertainty. The shaded vertical area is the dose interval for chow (grey) and HFD (blue) body weight adjusted STZ doses based on the mean (±SD) weight in our cohort. (F) Model predictions of dose-response curve showing fasting plasma glucose endpoint after 25 weeks, but with different timing of STZ dosing scheme at either week 1, 7 (our study), 14, 21.

Next, we used the validated model to predict the dose-response relationship between STZ and fasting plasma glucose. This was done by simulating several 25-week trials with a range of STZ dosages (0-10 mg total), with injections administered at week 7 over 4 consecutive days for both chow and HFD (Fig 5E). Here each dot represents the fasting plasma glucose endpoint (25 weeks) for a single simulated trial (the shaded area is the model uncertainty). The model predicts a sigmoidal relationship between the total STZ dose and endpoint plasma glucose, with maximum response of approximately 25 mM at higher doses. To illustrate the potential difference in outcomes between fixed body weight-adjusted dosing, we highlight the body weight adjusted-doses based on the mean body weight (± SD) at week 6 (shaded boxes Fig 5E). Here, focusing on HFD, we can observe that the body weight-adjusted dosing (based on mean body weight) resulted in different fasting plasma glucose values (15.5 – 21.2 mM) compared to the endpoint using fixed dose (17.6-19.1mM), based on data uncertainty captured by the model. Importantly, the model does not capture the full uncertainty in data (outliers), and the observation is based on the mean behaviour. Nevertheless, this may warrant further comparison between the two dosing schemes.

Lastly, we also investigated the effect on plasma glucose on timing of the STZ intervention. To do this we simulated four 25-week HFD interventions with the same STZ dosing schemes at different time points, either at weeks 1, 7, 14 and 21 (Fig. 5F). The model predicts that a larger total STZ dose, 5.76mg for dosing at 21 weeks compared to 4.95mg for 7 weeks, would be needed to reach the same endpoint fasting glucose (15mM).

## 4. Discussion

In this study, we characterized the effects of chow or HFD feeding combined with low or high STZ doses on multiple metabolic parameters over a 25-week period. We identified a robust protocol for inducing T2D in C57BL/6J mice, combining HFD feeding with anomer-equilibrated STZ administered at a fixed dose (mg) rather than body weight-adjusted (mg/kg). Historically, weight-adjusted dosing of STZ has been the standard approach for inducing diabetes in rodent models (26). However, studies comparing animals across differences in age, diet, sex, and species consistently report more severe glycaemic responses to STZ in heavier animals (11, 17, 26, 27). While these factors (age, diet, sex, and species) may each contribute to the variability in STZ sensitivity, an alternative explanation is that weight-adjusted dosing leads to relative overdosing in heavier animals, whereas fixed dosing (mg) may reduce this effect. Consistent with this, weight-adjusted dosing of hydrophilic compounds, such as STZ, is well recognised to increase the risk of overdosing in obese subjects (18, 19). Thus, fixed STZ dosing, especially in obese animals, may yield more consistent glycaemic responses. In support of this, HFD-STZ^low^ mice maintained obesity and insulin resistance while developing moderate hyperglycaemia and partial β-cell loss. Longitudinal assessment showed progression from HFD-induced insulin resistance with compensatory β-cell function to overt T2D following STZ-induced β-cell loss.

Considerable heterogeneity exists in protocols that combine HFD with STZ to induce T2D in mice, with wide variation in HFD duration, STZ dose and number of injections. Similar diabetic phenotypes (∼13–19 mM fasting glucose) have been achieved using markedly different strategies, including 3–12 weeks of HFD feeding followed by either a single high-dose or multiple low-dose STZ regimens (e.g. 1×100, 3×30, 3×40, 3×45, 3×75, or 5×50 mg/kg) (11, 28–32). This extensive heterogeneity highlights the need for a more standardized and reproducible protocol. Based on our findings, we propose using an anomer-equilibrated fixed STZ dose (4.8 mg total, administered as 1.2 mg per day over four consecutive days) in mice fed HFD for 7 weeks prior to STZ treatment. The duration of HFD-feeding is critical to establish a prediabetic state, with pronounced insulin resistance before β-cell loss is induced by STZ treatment, to better mimic the progression of T2D.

HOMA-β has been used as a surrogate index for monitoring β-cell function in studies using STZ (33, 34). However, HOMA-β was developed and validated for human physiology, and its use in animals requires caution and has been discouraged (25). When used, HOMA-β should be interpreted comparatively within a study, relative to a healthy control, and always in conjunction with HOMA-IR, as the index is inherently influenced by systemic insulin resistance (25). Despite the limitations of applying this index to rodent models, we observed a strong positive correlation between HOMA-β and pancreatic β-cell mass: HOMA-β explained 85% of the variance in insulin-positive β-cell area (R² = 0.85) across all diet/STZ combinations. Given that histological quantification of β-cell mass is labour intensive, HOMA-β provides a practical proxy for monitoring relative β-cell loss following STZ treatment. The β-cell loss observed in HFD-STZ^low^ mice closely mirrors that seen in humans at the time of T2D diagnosis (∼50% loss of β-cell function), further adding to the suitability of this model for studying effects of T2D *in vivo* (35). In contrast, human β-cell loss at T1D diagnosis is typically around 80%, though highly variable by age of diagnosis (36). Consistent with this, STZ^high^ mice (both chow and HFD) exhibited extensive (∼80-90%) β-cell loss, indicating that HFD-STZ^high^ mice have a metabolic profile that more closely models T1D than T2D.

An important strength of this study is the extended follow-up after STZ treatment, which allowed assessment of the stability of the diabetic phenotype over time. Over weeks 9–24, the HFD-STZ^low^ group showed remarkable consistency in weekly mean glucose (14.4–18.4 mM), insulin (1.35–1.69 ng/mL), HOMA-IR (28.0–35.7), and HOMA-β (58.9–96.0) (min–max range of weekly averages), indicating a sustained T2D-like state. This is notable given that other studies have reported a high remission rate of hyperglycaemia over longer follow-up periods, and it further suggests that fixed STZ dosing combined with α/β-anomer equilibration may reduce long-term variability in STZ-based T2D models (37).

Herein we also constructed a data-driven semi-mechanistic mathematical model capable of simultaneously describing STZ interventions (low and high doses) and vehicle (control) for chow and HFD mice. The model predictions of plasma glucose dynamics were also validated using independent-validation data from Gilbert *et al.* (11). There exist some previous modelling efforts of HFD and insulin resistance in mice. We have previously published a mathematical model of HFD-induced insulin resistance where we also explored β-cell dynamics (38). This model had a short-term focus (2 weeks) and did not include STZ intervention or β-cell data. Thus, our new mathematical model makes a valuable contribution both for simulating longer diets and also STZ interventions. Moreover, the current mathematical model has some limitations; the model agreement does not pass a formal χ^2^-test (when including all data simultaneously) on the residuals, but still fit qualitatively well to all data. Reasons for not passing a χ^2^-test is partially because we do not allow for any difference in parameter values (except for inputs) between cohorts, and we do not yet include mechanisms for age related insulin sensitivity. Still, the model could be improved in future work with a more mechanistic description of β-cell mass and insulin resistance progression, as well as extended with mathematical models for body-weight changes e.g. (39), to incorporate STZ effect on fat storage capabilities. The model predicted a sigmoidal dose–response relationship between STZ dose (mg) and fasting plasma glucose levels. A similar sigmoidal dose-response relationship has been observed in rats (40). The model prediction highlights that body weight adjusted dosing in HFD-fed mice may introduce high variance in fasting plasma glucose.

In summary, fixed-dose, anomer-equilibrated STZ administered after HFD feeding induced a stable T2D-like phenotype with low inter-individual variability over an extended follow-up period. This strategy may be particularly useful in HFD-fed mice, where variability in body weight can strongly influence conventional mg/kg dosing and contribute to inconsistent glycaemic outcomes.

## Supporting information

Raw data

## Acknowledgements

We thank the Lund Stem Cell Center Imaging Facility and Emanuela Monni for providing access to, and guidance in the use of, the Olympus VS-120 virtual slide microscope.

## Data and resource availability

All metabolic data generated and analysed in this study are provided in the Supplementary Excel file.

## Author contributions statement

Conceptualization: C.F., M.S.; Data curation: C.F., C.S.; Formal analysis: C.F., C.S., M.S., A.H.; Funding acquisition: C.F., M.S.; Investigation: C.F., C.S., A.H., J.M.; Methodology: C.F., M.S., C.S., A.H., J.M.; Project administration: C.F., M.S., K.G.S., C.S.; Software: C.S.; Supervision: C.F., M.S.; Visualization: C.F., C.S., M.S., J.M.; Writing – original draft: C.F., C.S., M.S.; Writing – review & editing: C.F., M.S., K.G.S., C.S., A.H., J.M.

## Use of AI tools

Large Language Models (LLMs) were used to support manuscript preparation by assisting with the initial drafting process and improving text for clarity, flow and readability.

## Supplementary

### Supplementary S1

**Supplementary Figure 1.**
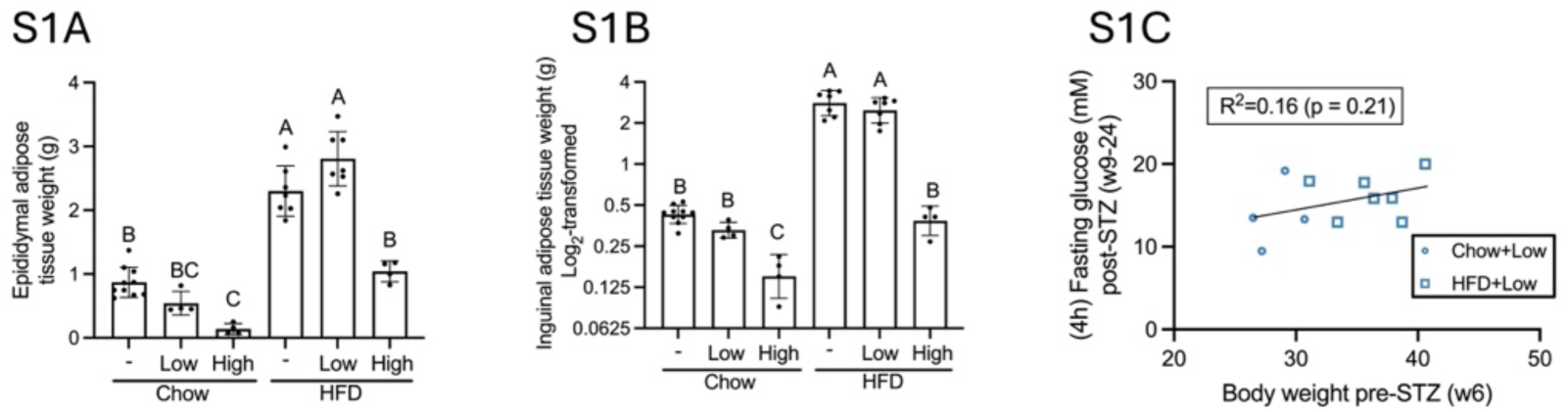
(A) Epididymal adipose tissue weight and (B) inguinal adipose tissue weight were measured at endpoint in chow- and HFD-fed mice receiving vehicle, STZ^low^, or STZ^high^. Inguinal adipose tissue weights are shown as log2-transformed values. Data were analysed by two-way ANOVA with diet and STZ treatment as factors, followed by Sidak’s multiple-comparisons test. Different letters indicate significant differences between groups based on Sidak’s post hoc multiple-comparisons test (p<0.05), groups sharing a letter are not significantly different. Data are displayed as mean ± SD. (C) Relationship between pre-STZ body weight at week 6 and post-STZ fasting glucose (mean weeks 9–24) in mice receiving a fixed dose of STZ. The analysis was performed to test whether mice with higher body weight at the time of injection showed a weaker glycaemic response to the same STZ dose. Linear regression showed no significant association between pre-STZ body weight and post-STZ fasting glucose (R2=0.16, slope= 0.23, 95% CI [-0.14 to 0.97], p=0.21).

### Supplementary S2

**Supplementary Figure 2.**
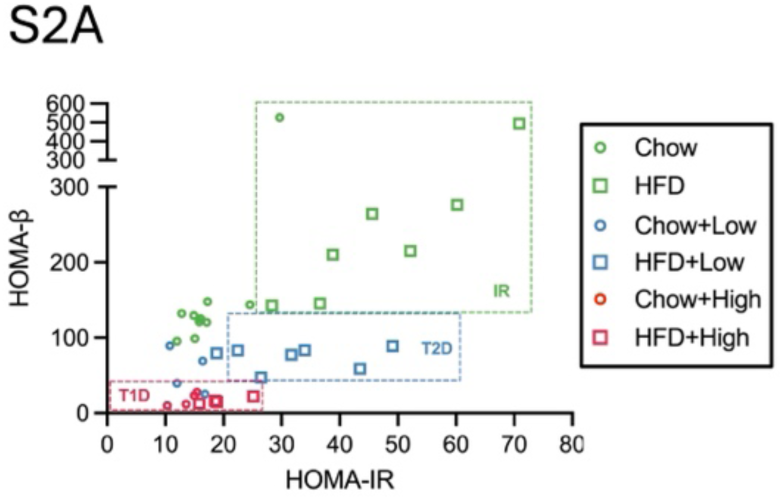
Relationship between HOMA-IR and HOMA-β (mean weeks 9-24) in chow- and HFD-fed mice receiving vehicle, STZ^low^, or STZ^high^. Outlined boxes indicate clustering of metabolic phenotypes: insulin resistance (high HOMA-IR, high HOMA-β) in HFD vehicle, T2D (high HOMA-IR, moderately reduced HOMA-β) in HFD-STZ^low^, and T1D (normal HOMA-IR, low HOMA-β) for chow-STZ^high^ and HFD-STZ^high^. Chow vehicle serves as a healthy control reference.

### Supplementary S3

#### Model description

This section will go through the mathematical model equation and all underlaying assumptions. The model analysis was done using MATLAB 2023b and the Systems Biology Toolbox (1). The model was developed using iterative hypothesis testing, with a multiride of candidate model structure being evaluated until one was found that could describe data sufficiently well.

The model was developed for the purpose of describing data for the levels of β-cells (A.U.), fasting plasma glucose (mM) and fasting plasma insulin (ng/mL) for controls, and STZ interventions in chow and HFD mice. To describe the Streptozotocin (STZ) intervention we implemented a one-compartment first-order kinetic pharmacokinetic model for describing the level of STZ in plasma (parameters: *k*_*clc_sty_*_, *V*_*stz*_ and *k*_*a_stz_*_). The parameter starting point where chosen based on the assumption of rapid plasma clearance of STZ. The state representing STZ in plasma was then allowed to permanently lower the levels β-cells in the model. The β-cells were represented by the following equation:

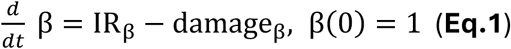

Where IR_β_ is a variable describing the increase of β cells during HFD, written as:

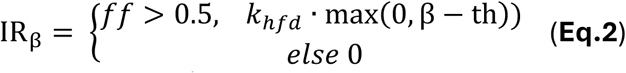

where *k*_*hfd*_(d^-1^) governs the increase in beta cells due to HFD, and *th* (a.u.) is the threshold for when enough beta cells are lost due to STZ to not have any diet-induced increase. The *damage* term was a hill equation:

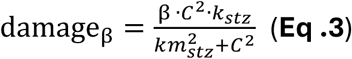

where C is plasma concentration of STZ, the maximal damage rate · *k*_*_stz_*_ and half-saturation constant *k*_*_stz_*_. To keep the model uncertainty lower we first optimized the STZ and the effect on β cells for all data and then kept the parameters governing STZ constant. To describe the effect on glucose-insulin interplay the model also included states representing fasting insulin and glucose. These state were loosely based on the glucose and insulin equation in the Topp et al. model (2). The fasting insulin was described as:

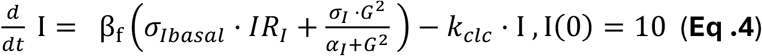

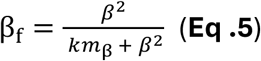

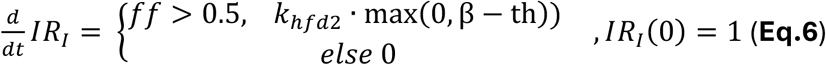

Where β_f_ is a variable descrbing the insulin producing capacity of the β cells, σ_*lbasal*_ the basal release of insulin, *IR*_*I*_ is a model state describing the change in this capacity during insulin resistance development, σ_*I*_ and α_*I*_ governs the glucose stimulated insulin release, and *k*_*clc*_ is the clearance of insulin. In the description of β_f_ (**E.q. 5**) the parameter *km_β_*, goverens the sensitivity between insulin secretion and beta cells.

In the description of *IR_*I*_*(**Eq. 6**) *k*_*hfd*2_ governs the diet-induced increase in insulin production capacity, and *th* is the same parameter from **Eq. 2**. Lastly, the model description of the fasting plasma glucose was described as:

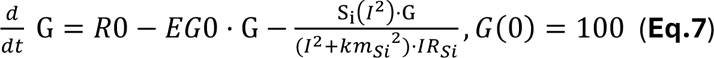

Where R0 is the basal release of glucose, *EG0* is the basal uptake of glucose, Si and *km_Si_* governed the insulin stimulated glucose uptake. Here, *IR_si_* is a model state representing the effect of increased insulin resistance, defined identically as Eq.6, but with a parameter **k*_*hfd*3_*.

#### Parameter values and parameter estimation

The parameter estimation was done using the MEIGO toolbox (3) using the *extend scatter search* (ESS) algorithm (with the dhc local solver option). To get a reasonable start guess we rescaled the Topp et al. model parameters using allometric scaling (from human to mice) and reproducing the same plasma glucose and insulin levels (even for smaller mice beta-cell mass). When running the parameter estimation, we ultimately had to allow for re-estimation of most parameters, however most parameters are still close to the re-scaled Topp et al. values. The following table has all estimated parameter values.

| Parameter Name | Unit | Estimated ($\theta_{best}$) |
| --- | --- | --- |
| $R_0$ | mg/dL/day | 7282 |
| $S_i$ | mL/(uU*day) | 378.691 |
| $EG_0$ | 1/day | 14.4945 |
| $\sigma_I$ | uU/(mL*day) | 0.0001 |
| $\alpha_I$ | (mg/dL)^2 | 18100 |
| $k_{clc}$ | 1/day | 208.21 |
| $Th$ | (a.u.) | 0.54 |
| $k_{hfd}$ | 1/day | 0.00458527 |
| $k_{stz}$ | (a.u.) | 591.72 |
| $km_{stz}$ | mg/kg | 1344.26 |
| $k_{hfd3}$ | 1/day | 0.102156 |
| $km_\beta$ | (a.u.) | 0.325071 |
| $km_{Si}$ | uU/mL | 113.881 |
| $\sigma_{Ibasal}$ | uU/(mL*day) | 7859.9 |
| $k_{hfd2}$ | 1/day | 0.0266882 |
| $k_{clcstz}$ | L/day | 0.02076 |
| $V_{stz}$ | L | 0.0175 |
| $k_{astz}$ | 1/day | 0.6 |

